# A single cell transcriptional atlas of early synovial joint development

**DOI:** 10.1101/2019.12.23.887208

**Authors:** Qin Bian, Yu-Hao Cheng, Jordan P Wilson, Dong Won Kim, Hong Wang, Seth Blackshaw, Patrick Cahan

## Abstract

Synovial joint development begins with the formation of the interzone, a region of condensed mesenchymal cells at the site of the prospective joint. Recently, lineage tracing strategies have revealed that Gdf5-lineage cells native to and from outside the interzone contribute to most, if not all, of the major joint components. However, there is limited knowledge of the specific transcriptional and signaling programs that regulate interzone formation and fate diversification of synovial joint constituents. To address this, we have performed single cell RNA-Seq analysis of 6,202 synovial joint progenitor cells from the developing murine knee joint from E12.5 to E15.5. By using a combination of computational analytics, *in situ* hybridization, and functional analysis of prospectively isolated populations, we have inferred the underlying transcriptional networks of the major developmental paths for joint progenitors. Our freely available single cell transcriptional atlas will serve as a resource for the community to uncover transcriptional programs and cell interactions that regulate synovial joint development.

## Introduction

Synovial joints are complex anatomical structures comprised of diverse tissues, including articular cartilage, synovium, fibrous capsule, and ligaments (Decker et al. 2014). Each of these tissues are susceptible to a range of diseases—both congenital and degenerative—and by common injuries that collectively have a profound global morbidity (den Hollander et al. 2019; Asahara et al. 2017). A better understanding of the inter- and intra-cellular networks that govern how these structures emerge during development will inform efforts to generate pluripotent stem cell derivatives for cell replacement therapy and disease modeling (Wang et al. 2019) and efforts to elicit regeneration *in situ* (Johnson et al. 2012). Moreover, an improved understanding of joint development will aid in identifying putative disease causing genes (Kelly et al. 2019).

Over the past two decades, lineage tracing has revealed much regarding the cell populations contributing to murine synovial joint development. It begins with the formation of the interzone (IZ), a region of condensed mesenchymal cells at the site of the prospective joint. In the mouse hindlimb, the IZ is initiated from a Col2a1^+^ Sox9^+^ pool of cells recruited from the mesenchymal condensation of the emerging limb bud starting at E11.5 (Hyde et al. 2008; Soeda et al. 2010). It is generally believed that chondrocytes at the presumptive joint de-differentiate (i.e. undergo a chondrocyte-to-mesenchymal transition) and begin to exhibit the flattened and layered morphology that is indicative of the IZ. A history of expressing Gdf5, a TGFβ ligand and critical contributor to joint formation (Storm and Kingsley 1999), marks cells that initially form the IZ or that later immigrate into it, and that subsequently go on to contribute to all of the major joint constituents including articular chondrocytes, ligament, meniscus, and synovium (Shwartz et al. 2016; Chen et al. 2016).

To gain a more comprehensive understanding of these developmental programs, bulk microarray expression profiling and RNA-Seq have been applied to the developing limb (Taher et al. 2011), to whole joints including the elbow and knee (Pazin et al. 2012), to the meniscus (Pazin et al. 2014), to the tendon (Liu et al. 2015), to connective tissue (Orgeur et al. 2018), and to laser-capture micro-dissected regions of the interzone (Jenner et al. 2014). While these investigations have yielded new insights into the genetic programs underpinning limb and joint morphogenesis, they provide limited resolution of the expression states for individual cell types due to the heterogenous nature of the samples profiled. With the advent of single cell profiling, it is now possible to detect transient populations of cells, to reconstruct developmental transcriptional programs, and to identity new cell populations (Guo et al. 2010; Kumar et al. 2017). For example, Feng et al revealed molecular signatures and lineage trajectories of an interzone related Lgr5^+^ population in the murine E14.5 knee joint that contributes to the formation of cruciate ligaments, synovial membrane, and articular chondrocytes (Feng et al. 2019).

Here, we applied single-cell RNA-sequencing on Gdf5-lineage cells of the murine hindlimb to determine the transcriptional programs of early synovial joint development. In contrast to the recent study of Feng et al 2019, which focused on lineage divergence of a specific Lrg5^+^ interzone population, we sought to characterize formation of the entire IZ and to discover the extent to which heterogeneity in the nascent interzone is resolved into the distinct lineages that are apparent later at cavitation. Therefore, we sequenced Gdf5-lineage cells from the presumptive joint of the hindlimb from E12.5 (prior to frank IZ formation) through E15.5 (coinciding with cavitation). We combined computational analytics and *in situ* hybridization to infer the lineage relationships of joint progenitors and to identify the combinatorial transcriptional programs that mediate the elaboration of the interzone into the major synovial joint lineages. We found that early Gdf5-lineage enriched cells consist of sub-populations with chondrogenic or fibrous-lineage bias. Furthermore, we discovered within the chondrogenic-biased population were two distinct sub-populations that followed similar trajectories to de-differentiate into IZ cells, supporting a model of regionally and temporally complex IZ specification (Shwartz et al. 2016). To aid the community in discovering additional transcriptional programs and in inferring cell interactions that contribute to synovial joint development, we have made this data freely and easily accessible with a web application at http://www.cahanlab.org/resources/joint_ontogeny.

## Results

### Gdf5Cre^+^ cells in the hind limb from E12.5 to E15.5 Gdf5^Cre^R26^EYFP^ mice are primarily located in the interzone, articular cartilage, ligament, menisci, and synovium, as well as in other non-joint tissues

Gdf5-lineage cells contribute to several components of synovial joint, including articular cartilage, meniscus, ligaments, and synovium. To isolate joint progenitors, we crossed Gdf5 promoter driven Cre mice with the R26 reporter mice in which loxP-flanked STOP sequence followed by the EYFP inserted into the Gt(ROSA)26Sor locus, allowing us to identify Gdf5-lineage cells by YFP expression. We used fluorescent immunohistochemistry to determine the spatial and temporal pattern of YFP. At E12.5, YFP is mainly expressed in the presumptive joint area including part of the bone anlagen and the surrounding mesenchyme (**Fig 1**). At E13.5, YFP^+^ cells are more centered in the interzone (IZ) and in the surrounding connective tissue; they are sparse in the anlagen of the femur and tibia. By E14.5, YFP^+^ staining is mainly present at the area of future articular cartilage (AC), synovium and surrounding soft tissue. YFP expression becomes obvious in menisci one day later. YFP^+^ cells are also seen in AC, epiphyseal cartilage, and synovium at E15.5.

**Figure 1:**
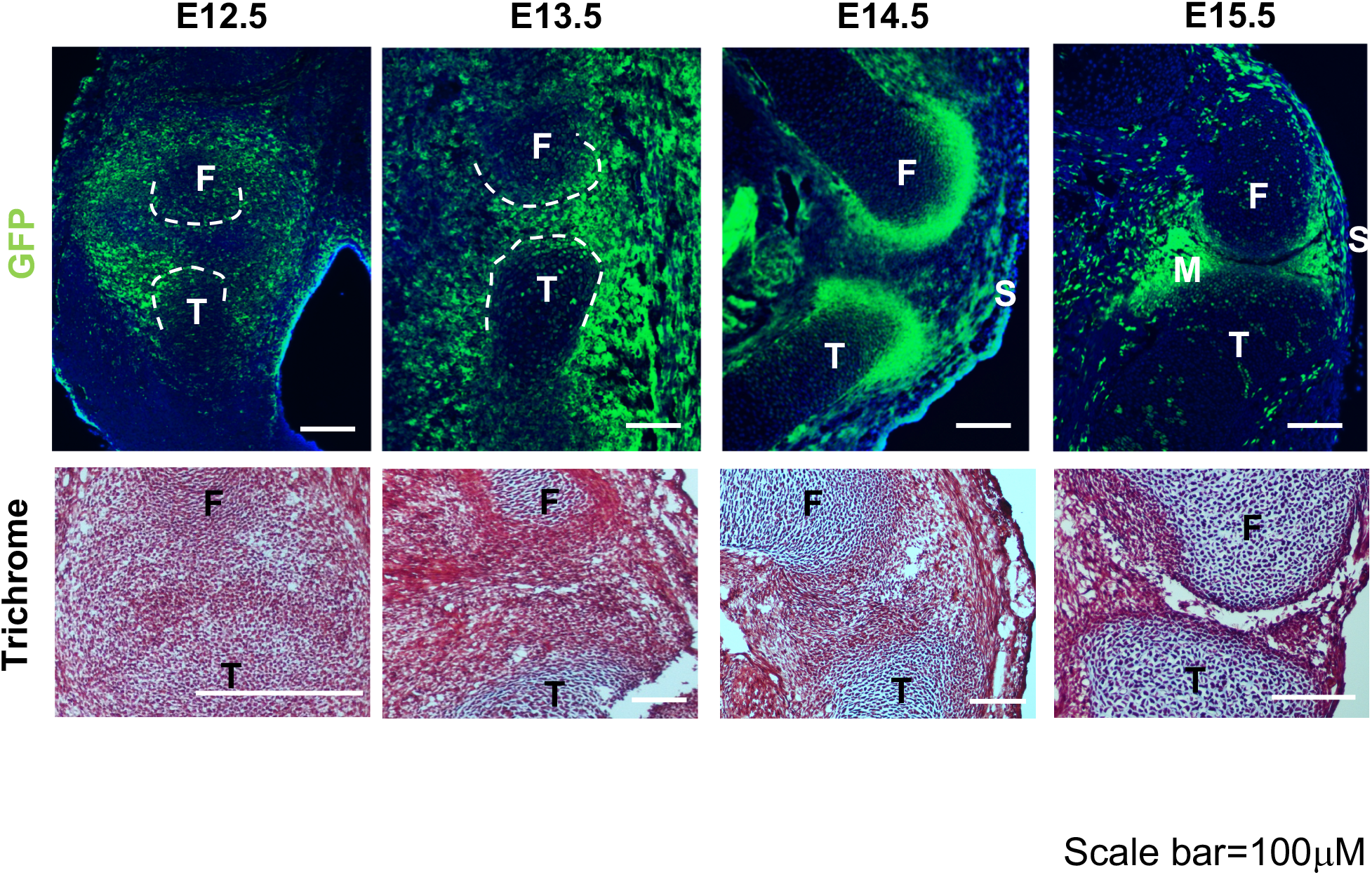
Top: localization of Gdf5-lineage cells in murine hindlimb. Bottom: Cell density and morphology during joint formation as shown by Trichrome staining. Scale bar = 100.

We observed “ectopic” YFP expression in non-joint tissues such as the dermis and muscle, consistent with prior reports (Roelofs et al. 2017). However, because our scRNA-Seq analysis pipeline includes a “cell typing” step (see below), we were able to identify these non-joint cells *in silico* and exclude them from our in-depth analyses that focus on the Gdf5-lineages of the joint. We refer to cells that passed our *in silico* filtering as Gdf5-lineage enriched (GLE) cells rather than YFP^+^ Gdf5-lineage cells because we cannot absolutely prove that YFP expression tracks with Gdf5-lineage in this system. Nonetheless, our staining combined with prior reports examining Gdf5cre cells in the limb, indicate that GLE cells are major cellular contributors to the knee joint. Therefore, determining their transcriptomes will yield insights into the genetic circuitry that accompanies IZ formation and the emergence of articular components such as ligament and tendon.

### GLE cells form three distinct super clusters across two major developmental stages

To define the transcriptional states of joint cells and their progenitors during landmark developmental events, we isolated YFP^+^ cells from the hind limbs of male embryos from E12.5 (the time just prior to frank IZ formation) to E15.5 (before cavitation). To minimize contamination with Gdf5-lineage cells from the ankle and digits, we manually dissected the region of the limb containing the presumptive joint and excluded the paw (**Supp Fig 1A**). Then, we collected Gdf5 lineage cells by fluorescence activated cell sorting (FACs) of YFP^+^ cells after enzymatically disassociating the presumptive knee joint region (**Supp Fig 1B**). We loaded approximately 6,000 cells for single cell RNA-Seq library preparation using the 10x Genomics platform, and sequenced the transcriptome of ~1,000 to 5,000 cells at a target depth of 100,000 reads per cell (**Table 1**).

**Tables 1:** Statistics on cells collected for scRNA-Seq. ‘Cells captured’ was determined by 10X CellRanger. GLE cells indicate the number of cells remaining after excluding cells unlikely to be GDF5-lineage, including immune cells, neural crest cells, and endothelial cells.

| Stage | YFP<sup>+</sup><br>cells | YFP<sup>+</sup><br>cells % | cells captured | GLE cells | Reads | Reads/cell | Genes/cell |
| --- | --- | --- | --- | --- | --- | --- | --- |
| E12.5 | 74.8K | 35.59% | 3,107 | 1964 | 489,942,625 | 157,689 | 3,403 |
| E13.5 | 113.9K | 11.26% | 4,786 | 2433 | 347,861,418 | 72,683 | 1,755 |
| E14.5 | 54.1K | 3.7% | 1,888 | 993 | 359,671,420 | 190,503 | 2,835 |
| E15.5 | 50.0K | 7.57% | 1,099 | 812 | 365,318,792 | 332,410 | 2,835 |

After performing quality control to remove potential doublets and low-quality libraries, we sought to identify the major transcriptional states in our data by clustering using the Leiden graphbased community detection algorithm (Traag et al. 2019). We found 14 clusters, many of which contained cells from multiple timepoints (**Supp Fig 2A**). To determine the cell type of each cluster, we used SingleCellNet to classify individual cells based on a well-annotated reference data set (Tan and Cahan 2019), and we used differential gene expression to identify marker genes of cell types that are not included in current single cell reference data sets (e.g. neural crest cells and melanocytes). This approach identified eight clusters made up of non-joint cell types including myoblasts, immune and red blood cells, neural crest cells and melanocytes, and endothelial cells (**Supp Fig 2B**). After removing these non-joint cells, we re-clustered the data and we performed differential gene expression analysis (**Supp Fig 2C**). All clusters had detectable levels of the osteochondral transcription factor (TF) Sox9 except one, which had high levels of genes associated with dermis, including Twist2 and Irx1 (**Supp Fig 2D**). To localize the cells in this cluster, we performed *in situ* hybridization (ISH), confirming that they are dermal cells (**Supp Fig 2E**), and we excluded these cells from further analysis. Finally, we performed cell cycle analysis by scoring each cell according to its likely phase (G1, G2M, or S) based on expression of canonical cell cycle-related genes (**Supp Fig 2C**). We removed two clusters, which were comprised predominately of cells in G2M or S phase, as we found that including these cells confounded downstream analysis. This cell trimming process resulted in a data set of 6,202 synovial joint GLE cells.

Next, we asked whether there were discernible transcriptional profiles that spanned timepoints. To address this question, we clustered all of the GLE cells and uncovered three ‘super-clusters’ (SCs), two of which contain a plurality of cells from more than a single timepoint (**Fig 2A-B**). One of the clusters corresponds roughly to developmental time: SC1 is 98.2% E12.5 cells. The other two SCs are mixtures, with SC2 and SC3 predominately made up of cells from E13.5-E15.5. To gain a better understanding of these SCs, we examined the expression of genes with well-established roles in limb and joint development. Prrx1 and Pitx1 are preferentially expressed in the early SC1 (**Fig 2C**), consistent with their roles in specifying limb mesenchymal cells from lateral plate mesoderm (Bobick and Cobb 2012; Marcil et al. 2003; Wang et al. 2018). Shox2, regulating onset of early chondrogenesis (Bobick and Cobb 2012) has a similar expression pattern. Since many cells in SC1 express Sox9 but few express Col2a1, it is likely that this supercluster is comprised of a mixture of progenitor cells of mesenchymal character and chondroprogenitors. 25% of SC1 cells express the IZ marker Gdf5, and thus may represent de-differentiated chondrocytes. SC2 is similar to SC1 in expression profile, but it also preferentially expresses Sox9, Gdf5, Col11a1 and Col2a1, suggesting that this SC is likely to contain a mixture of IZ cells and transient chondrocytes (Zhao et al. 1997). SC3 cells express fibrous related genes Col3a1, Col1a1, Lgals1 (Dasuri et al. 2004), Dcn (Havis et al. 2014), indicating that SC3 largely consists of fibroblast-related cells.

**Figure 2:**
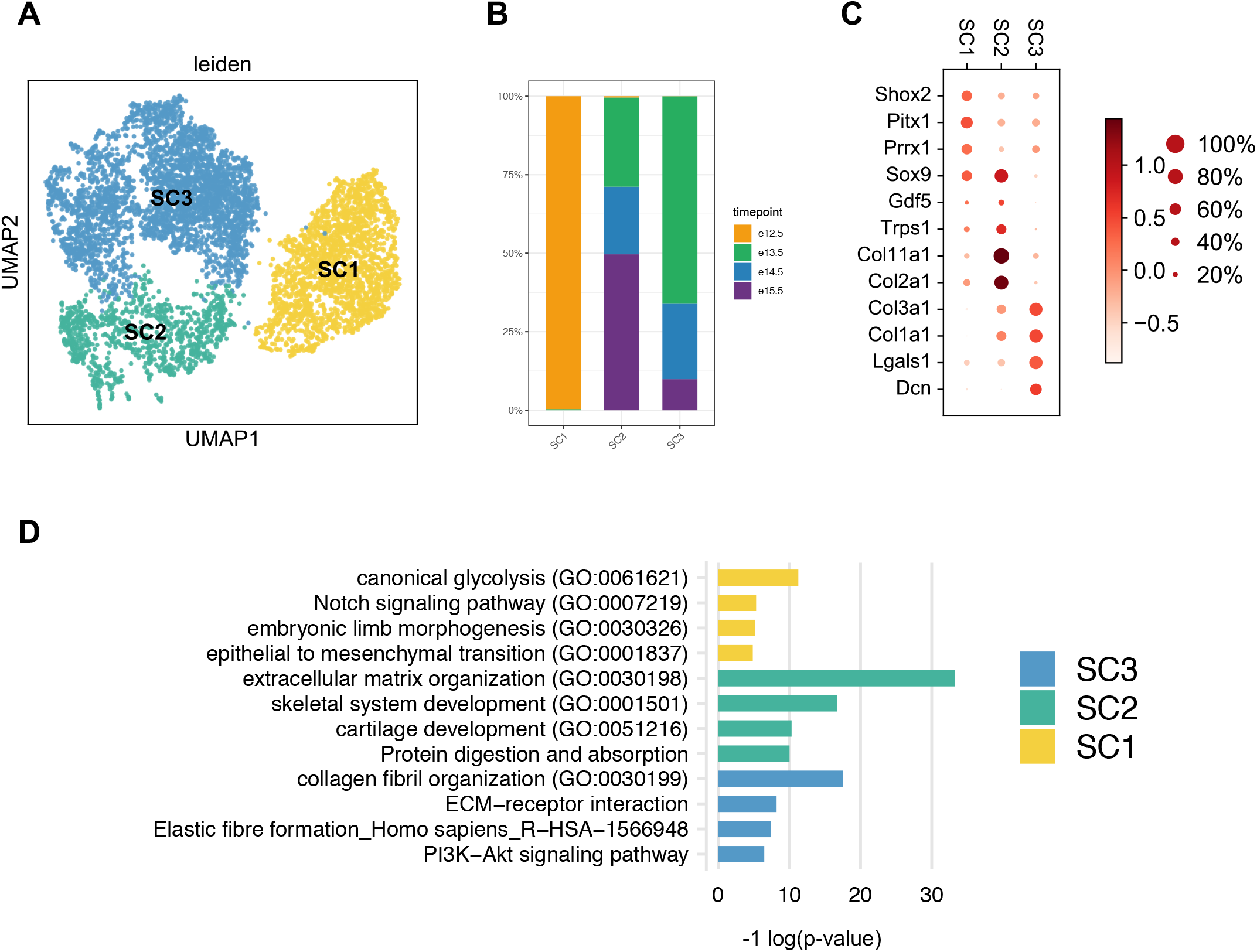
scRNA-Seq of Gdf5-lineage enriched cells during knee development. Leiden clustering and UMAP embedding of the five distinct superclusters of GLE cells (A). The proportion of cells from each timepoint varies across superclusters (B). Expression of genes well-characterized in limb and joint development (C). Size of each dot reflects the percent of cells in which the gene is detected within the supercluster. The color indicates mean expression, including cells in which there is no detectable expression. (D) Supercluster gene set enrichment analysis, showing selected categories. Complete results are in Supplemental Table 1.

Gene set enrichment analysis largely corroborated our supervised annotation of the superclusters (**Fig 2D**). SC1 is enriched in limb and joint development-associated pathways including embryonic limb morphogenesis, Notch signaling (Jiang et al. 1998), and epithelial to mesenchymal transition. SC2 is enriched in extracellular matrix (ECM) organization, skeletal system development, and cartilage development. SC3 involvement in fibrous differentiation is supported by the enrichment of collagen fibril organization and elastic fiber formation.

Taken together, this analysis has revealed three major transcriptional states of GLE cells in synovial joint development. It has also hinted at substantial heterogeneity within SCs. To more clearly define the cell types and states of GLE cells, we next analyzed each SC separately, as described in the following sections.

### Two categories of early GLE cells: chondrogenic and mesenchymal

By applying Leiden clustering to only SC1, we identified two sub-clusters: SC1_A and SC1_B (**Fig 3A**). SC1_A has high expression levels of genes associated with chondrogenesis (e.g. Sox9 and Col2a1) and the IZ (e.g. Nog and Gdf5) (Ray et al. 2015; Hartmann and Tabin 2001; Storm and Kingsley 1996). SC1_B exhibited high expression levels of genes associated with fibrous and mesenchymal cells such as Col3a1 and Col1a2 (Niederreither et al. 1992), as well Osr1, which is mainly expressed in the outer mesenchyme (**Fig 3B**) where it promotes fibroblast differentiation and inhibits chondrogenesis (Stricker et al. 2012). These results suggest that SC1 is comprised of chondroprogenitors and early chondrocytes of the limb anlagen, nascent IZ cells, as well as the non-chondrogenic mesenchymal cells situated outside the anlagen. We tested and confirmed this conjecture using ISH for genes indicative of each cluster (**Fig 3C-D**).

**Figure 3:**
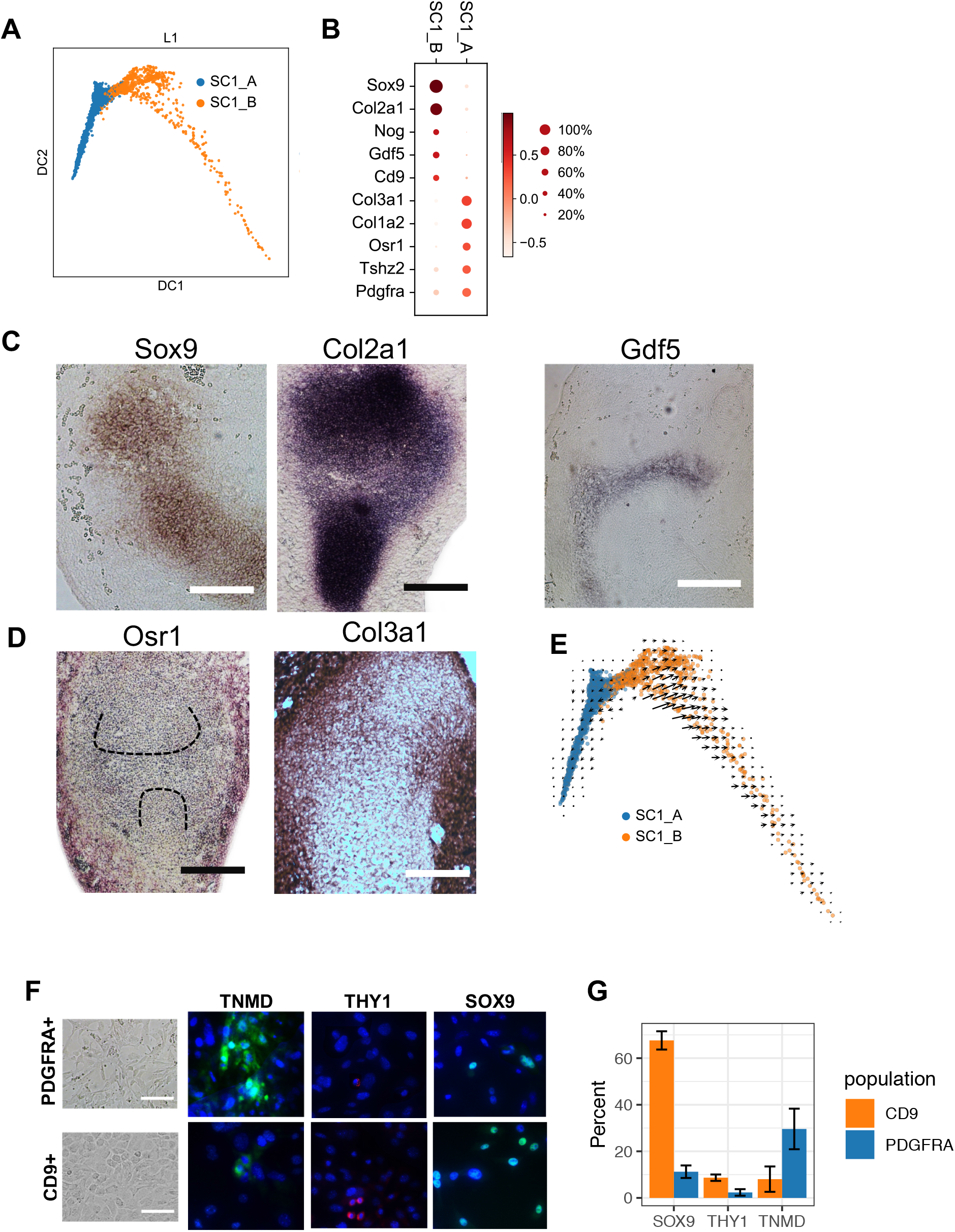
SC1 is composed of chondrogenic and mesenchymal fated cells. (A) Leiden clustering and diffusion map embedding SC1. (B) Dot plot expression of representative genes differentially expressed between SC1_A and SC1_B. (C) ISH detection for SC1_A and SC1_B representative genes. (E) RNA Velocity analysis. Arrows indicate the predicted future state of SC1 cells, showing a minimal transition between SC1_A and SC1_B. (F) In vitro culture of YFP^+^/Pdgfra^+^ and YFP^+^/Cd9^+^ hindlimb cells from e12.5 embryos shows distinct morphology of the cells (left). Immunofluorescence staining of tendon and ligament marker TNMD, fibroblast marker THY1, and chondrocyte regulator SOX9 (right). (G) Quantification of the proportion of cells positive for each marker.

To determine the lineage relationship between these clusters we performed RNA Velocity analysis (La Manno et al. 2018). Our results predicted that there is little-to-no transition between SC1_A and SC1_B (**Fig 3E**). To test this prediction, we prospectively isolated E12.5 YFP^+^ cells using antibodies specific for SC1_A (CD9) or SC1_B (PDGFRA), and measured lineage specific maker expression after culturing the cells *in vitro* for seven days. Cells from the PDGFRA^+^ population exhibited a mesenchymal morphology, whereas cells from the CD9^+^/PDGFRA^-^ population exhibited a chondrocyte-like morphology (**Fig 3F, left**). Consistent with their respective shapes and appearances, the PDGFRA^+^ population yielded a substantially higher proportion cells positive for the tendon and ligament marker TNMD compared to the CD9^+^/PDGFRA^-^ population, and a lower proportion of cells positive for the chondrogenesis regulator SOX9 as measured by immunofluorescence (**Fig 3F, right and Fig 3G**). While the CD9^+^ population yielded more THY1-positive cells, neither group had a substantial fraction of positive cells. The fact that both populations were not mutually exclusive for TNMD and SOX9 expression can be explained by incomplete lineage commitment, by the imperfect ability of PDGFRA to mark SC1_A and of CD9 to mark SC1_B, and by impurity in the FACS gating. With these caveats in mind, the data do support a model where the *in vitro* differentiation propensity of SC1_A is towards a tenocyte/ligamentocyte fate, whereas the *in vitro* propensity of SC1_B is towards a chondrocyte fate.

### Diverse origins of nascent interzone

While most SC1_B cells expressed Sox9, we noticed that they were heterogeneous in terms of IZ- and chondrocyte-related genes, suggesting that this cluster consisted of sub-populations or sub-states. To examine this further, we clustered SC1_B alone and identified four clusters: SC1_B1 to SC1_B4 (**Fig 4A**). SC1_B4 was marked by high levels of Col2a1 and Matn1, indicating that it contained cells destined to become transient chondrocytes (Hyde et al. 2007) (**Fig 4B**). The three other clusters expressed both chondroprogenitor transcription factors (e.g. Sox5, Sox6, and Sox9), as well as the IZ marker Gdf5. These clusters varied in the extent to which they expressed other IZ-related genes: SC1_B3 had high levels of Sfrp2, Vcan, and Trps1, whereas SC1_B2 had the highest level of Ebf1, Jun respectively (**Fig 4B**) (Choocheep et al. 2010) (Norris et al. 2007; Choocheep et al. 2010; Kunath et al. 2002; Salva and Merrill 2017).

**Figure 4:**
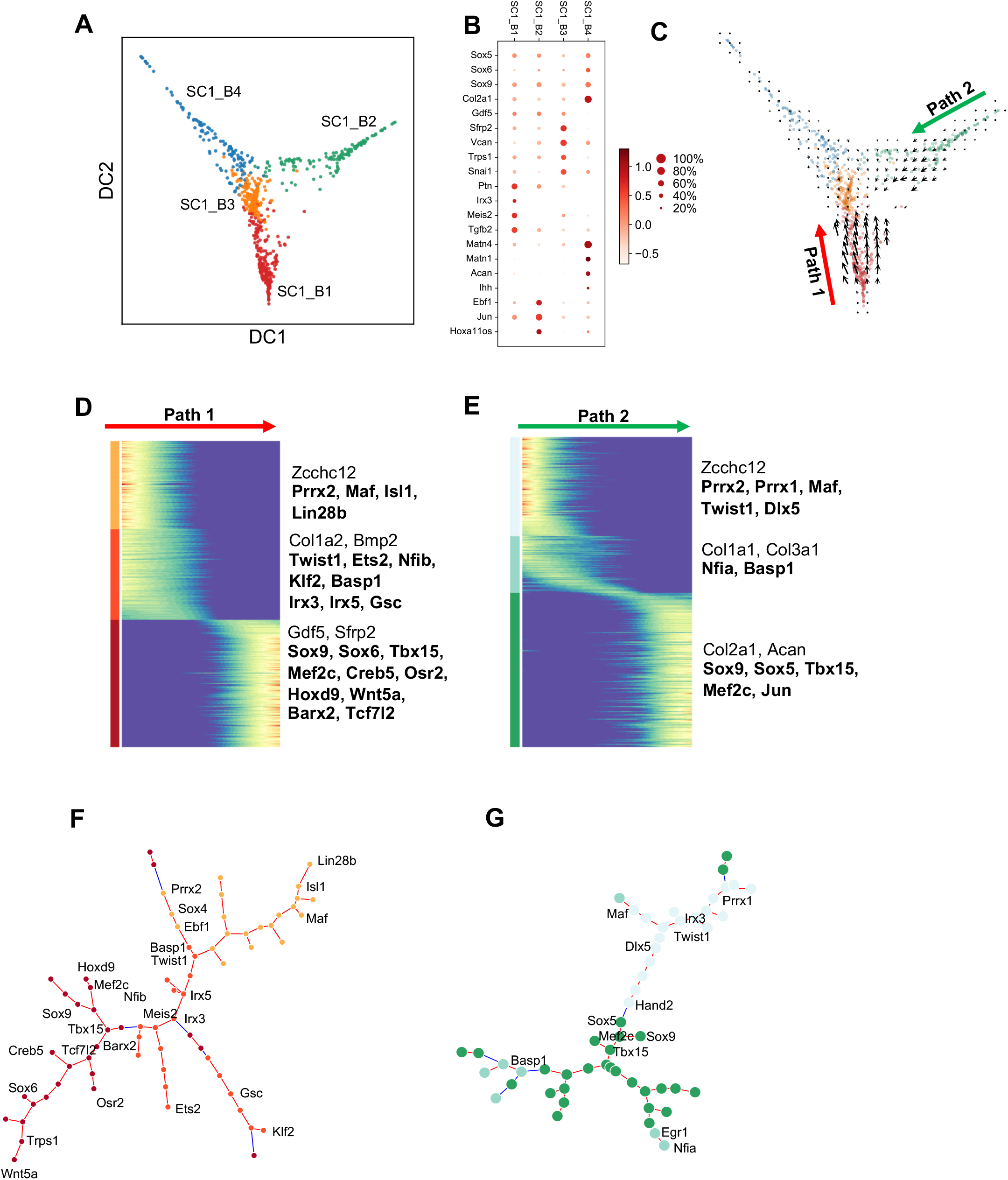
Two SC1_B sub-populations converge to a common interzone-like state. (A) Leiden clustering and diffusion map embedding of SC1B. (B) Dot plot of expression of representative genes differentially expressed between SC1B sub-clusters. (C) RNA velocity indicates converging trajectories of SC1_ B1 and SC1_ B2. Epoch analysis identifies three Epochs of gene expression in Path1(D) and Path 2 (E). Selected genes from each Epoch are listed on right, and Epoch identified regulators are in bold. (F-G) Minimal spanning tree representation of Epoch-reconstructed gene regulatory network.

With the exception of SC1_B4, we hypothesized that these clusters represented distinct stages of IZ formation. To explore this hypothesis, we performed RNA velocity analysis. Consistent with the notion that SC1_B4 consists of transient chondrocytes, the trajectories of the other, presumptive IZ, clusters did not lead to it (**Fig 4C**). Rather, the only trajectories were an apparent convergence of IZ clusters SC1_B1 and SC1_B2 to at a singular IZ expression state defined by high expression of IZ markers such as Sfrp2 (Pazin et al. 2012) and Vcan (Choocheep et al. 2010) in SC1_B3. To better understand the regulatory networks that contributed to this convergence, we subjected these clusters to Epoch analysis (manuscript in preparation). In brief, this tool takes as input pseudotime-ordered scRNA-Seq data. Then it identifies temporally regulated genes and periods of gene activity, reconstructs gene regulatory networks in a temporally sensitivity manner, and it proposes candidate regulators of transitions between expression states. To use this tool, we first ordered the cells along a pseudotemporal axis as defined by diffusion-based pseudotime (Haghverdi et al. 2016), with two roots, or starting points, selected based on the RNA velocity analysis. Then, we used Epoch to identify genes temporally regulated along each of these converging trajectories, or paths. Each path held three classes of genes: those with expression that peaked at early, in the middle, or later in the trajectory (**Fig 4D-E**). Path 1, which is defined by cells from SC1_ B1, starts with an expression of limb mesenchyme (high Prrx2 (Leussink et al. 1995), Zcchc12 (Li et al. 2009)), then expression of Col1a2 and Bmp2 peaks in the middle stage, and it ends in the high Gdf5 and high Sfrp2 state. Epoch predicted that the major regulators of the first stage are Maf, a known regulator of chondrocyte differentiation (MacLean et al. 2003), Isl1 (Yang et al. 2006), Ebf1 and Sox4, detected in IZ with unknown mechanism (Jenner et al. 2014; Bhattaram et al. 2014) and Lin28b, an indicator of embryonic to adult transitioning (Zhang et al. 2016) (**Fig 4D**).

The middle stage of Path 1 was predicted to be regulated by epithelial to mesenchymal transition regulator Twist1(Liu et al. 2017), and chondrogenic regulators Klf2 (Cameron et al. 2009) and Ets2 (Karsenty and Wagner 2002). Other regulators included Meis2, which was previously reported as expressed in the knee IZ (Pazin et al. 2012) and Nfib, a homolog of Nfia which maintains the IZ domain (Singh et al. 2018).

The later stage of Path1 is predicted to be regulated by Sox9, which is considered to decrease in expression during IZ formation (Soeda et al. 2010); IZ morphogenesis regulator: Sox6 (Dy et al. 2010); Tcf7l2, which mediates crosstalk the between Hedgehog and Wnt signaling that promotes IZ differentiation (Rockel et al. 2016). Epoch also identified Osr2, Barx2, Hoxd9, Wnt5a, and Trps1 as important contributors to the late stage of Path 1. Many of these factors have previously been reported to be associated with IZ: Osr2 contributes to IZ expression of Gdf5 (Gao et al. 2011); Barx2 is upregulated in the presumptive IZ (Meech et al. 2005); Hoxd9 regulates sesamoids formation from IZ (Khoa et al. 1999; Fromental-Ramain et al. 1996); Wnt5a was detected in digital IZ and is downregulated at cavitation (Church et al. 2002); Trps1 acts downstream of Gdf5 to promotes chondrogenesis (Itoh et al. 2008). Path 2, which is defined by cells from SC1_ B2, also starts with an expression state of mixed limb mesenchyme (high Prrx1,2 and Zcchc12), but a distinct set of IZ related TFs including Dlx5 (Ferrari and Kosher 2006) and Hand2 (Askary et al. 2015) are involved in regulating early stage of transition. Many of the regulators and target genes of the middle and tertiary stages Path 2 are shared with Path 1 (**Fig 4D-G**). For example, the middle stage of Path 2 is marked by peak expression of Col1a1, Nfia, and Basp1. Similarly, the final stages of both paths are marked by peak expression of Sox9, Mef2c, Tbx15. However, a notable difference in the paths is in the early stages where Path 1 is regulated by Irx3, Irx5 and Meis2, which are preferentially expressed in the proximal anterior portion of the developing limb (Li et al. 2014) (Capdevila et al. 1999). This suggests that the SC1_B1 and SC1_B2 start at distinct states reflecting remnants of spatial patterning of the condensing mesenchymal cells of the limb. However, as they differentiate, they leverage the same, or highly similar GRNs, to converge on the IZ state. Overall, the trajectories presented here indicate a process in which similar but separately patterned early chondroprogenitors follow parallel paths, with shared landmarks (e.g. an intermediate stage in which IZ regulators Sox9 peaks), before reaching an IZ-like state. All of the regulators predicted by Epoch are listed in **Supp Table 1, 2**.

### IZ formation

Compared to SC1, many SC2 cells had high levels of more IZ-related genes such as Cd44 and Sfrp2; other SC2 cells exhibited more established chondrocyte profiles. To resolve this population heterogeneity, we performed clustering on SC2 and identified two groups (**Fig 5A**). SC2_A was enriched in chondrocyte-related genes Col2a1, Col9a1, Sox9 (**Fig 5B**). We confirmed the expression co-localization of these genes in the anlagen by ISH (**Fig 5C**). SC2_B, on the other hand, exhibited IZ features based on higher expression of Gdf5, Sfrp2, and Col3a1 (**Fig 5B**). We confirmed the IZ localization of these genes by ISH (**Fig 5D**). To understand the lineage relationship between these clusters, we performed RNA Velocity analysis, finding that approximately half of the SC2_A cells were transitioning to SC2_B (**Fig 5E**), suggesting that GLE anlagen prechondrocytes continue to de-differentiate and contribute to the IZ.

**Figure 5:**
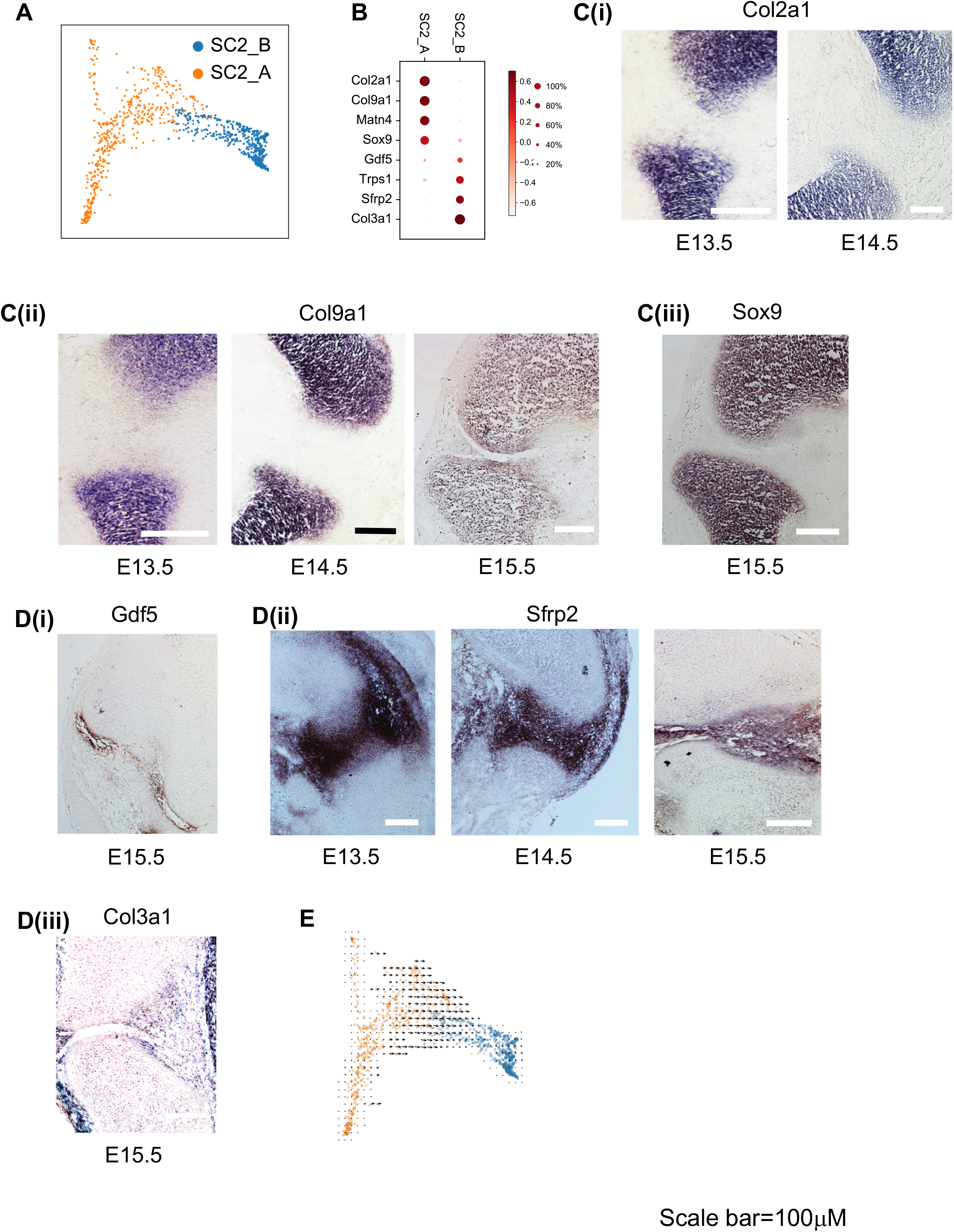
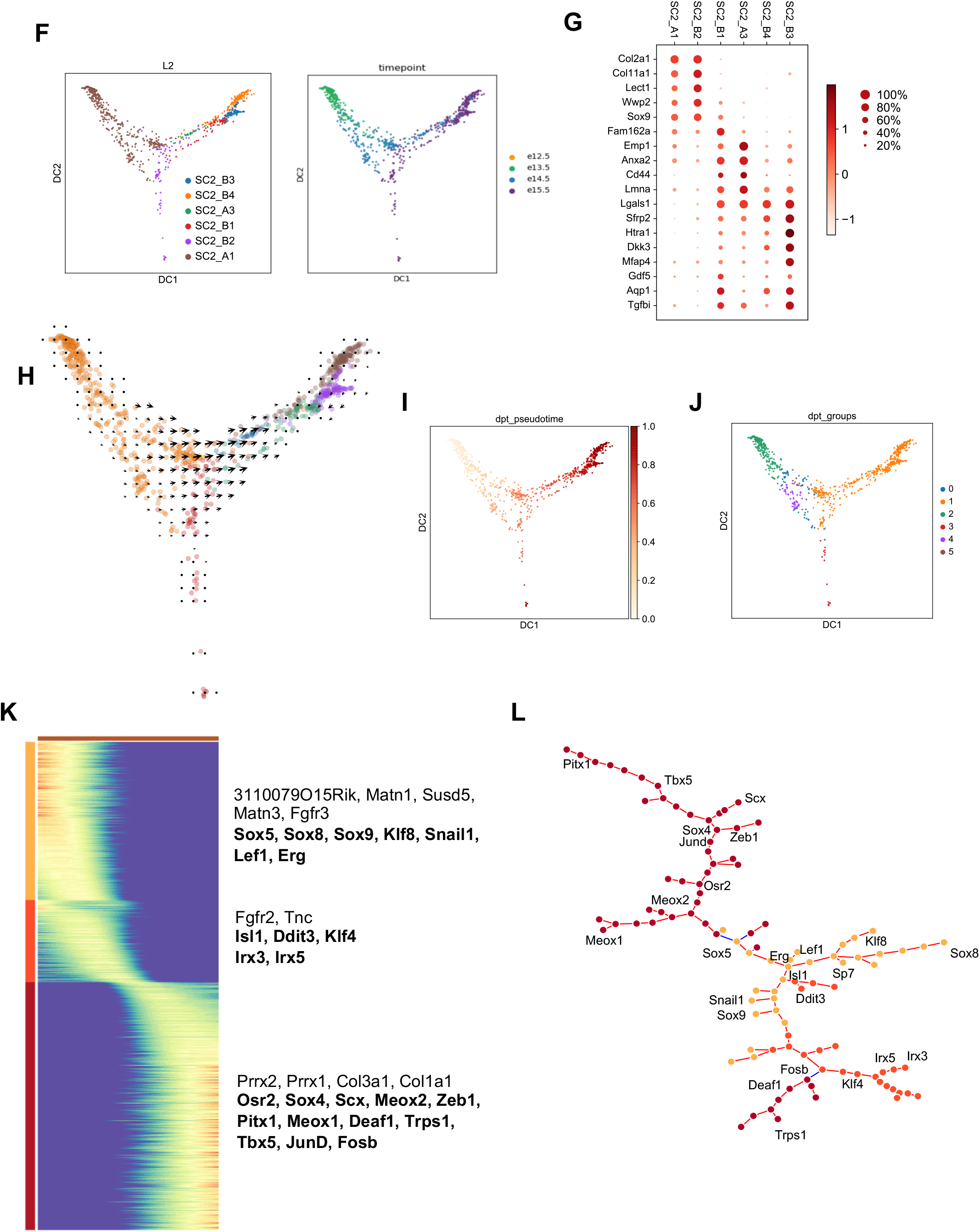
SC2 is composed of interzone fated cells. (A) Leiden clustering and diffusion map embedding SC2. (B) Dot plot expression of representative genes differentially expressed between SC2_A and SC2_B. (C,D) ISH detection for SC2_A and SC2_B representative genes. (E) RNA Velocity analysis. Arrows indicate the predicted future state of SC2 cells, showing a minimal transition between SC2_A and SC2_B. **IZ formation is a chondrocyte to mesenchymal cell transition process.** (F) Leiden clustering and diffusion map embedding of SC2, colored by subgroups or timepoints. (G) Dot plot of expression of representative genes differentially expressed between SC2 sub-clusters. (H) RNA velocity indicates a developmental path connecting sub-clusters. (I, J) Leiden clustering of SC2, colored by pseudotime (I) and groups (J). (L) Epoch analysis identifies three Epochs of gene expression based on group1 (orange) shown in (J). Selected genes from each Epoch are listed on right, and Epoch identified regulators are in bold. (M) Minimal spanning tree representation of Epoch-reconstructed gene regulatory network.

The fact that a substantial fraction of the SC2_A cells were not transitioning to IZ suggested population substructure. To explore this, we performed clustering on each of SC2_A and SC2_B, finding three and four subsets respectively. One of the SC2_A sub-clusters, SC2_A2, consisting of about 20 cells, expressed Ihh and Cd200, suggestive of a pre-hypertrophic state (**Supp Fig 3A**). As these cells were not predicted to be related to any of the other clusters by RNA velocity, we excluded them from further analyses (**Fig 5F**). We found that SC2_A1 and SC2_B2 have similar expression patterns in chondrogenic genes Col2a1, Col11a1, Sox9, Wwp2 (Zhao et al. 1997; Hyde et al. 2007; Akiyama and Lefebvre 2011) (**Fig 5G**). The other four clusters have lost expression of Col2a1 and have upregulated expression of IZ-related genes including Gdf5, Cd44 (Hartmann and Tabin 2001), Sfrp2, Htra1(Oka et al. 2004), and Dkk3 (Witte et al. 2009) (**Fig 5G**). Repeating RNA velocity on these clusters recapitulated the results of a transition from chondrocyte to IZ state when applied to all SC2 cells (**Fig 5H**).

To identify the GRN contributing to this transition, we applied Epoch to the group of cells that exhibited a concerted velocity from chondrogenic to IZ-like, as defined and ordered by diffusion based pseudotime (**Fig 5H,I,J**). Epoch analysis revealed different early regulators as compared to the programs identified in IZ initiation at E12.5 (**Fig 5K, Supp Table 3**). Here, in the early part of the path, some of the IZ progenitors appear to be outer IZ based on the higher level of expression of outer IZ related genes: 3110079O15Rik, Matn1, Susd5, Matn3, Fgfr3 and coexpression of Gdf5 (**Supp Fig 3B**). Epoch predicted that the major regulators of the first stage included the known IZ regulator Erg (Iwamoto et al. 2007); Lef1, an effector of canonical Wnt signaling with multiple roles in early IZ specification (Guo et al. 2004); Klf8 (Wang et al. 2011) and Snail1, two epithelial to mesenchymal transition (EMT) inducers, indicating a shared regulatory program between chondrocyte-to-mesenchymal transition and EMT (Vincent et al. 2009; Lin et al. 2014) (**Fig 5L**). The middle stage is marked by upregulation of both fibrogenic genes such as Fgfr2, Ddit3 (Caterson and Melrose 2018) and chondrogenic genes such as Isl1(Yang et al. 2006), Tnc (Grogan et al. 2013). This suggests that the middle stage is a transition state in which cells exhibit properties of both chondrocyte and IZ cell. Isl1 and Ddit3 also act as the predicted regulators for middle stage, as does Klf4, which promotes the expression of Col1a1 that is required for IZ morphogenesis (Orgeur et al. 2018). Interestingly, the mesenchyme markers Prrx2, Prrx1, Col3a1, Col1a1 expression turn back to the peak level at late stage, indicating there is a dedifferentiation of chondrocytes to mesenchymal cells in IZ development. The major regulators of the late stage include: Scx, an inducer of ligament/tendon differentiation (Anderson et al. 2006) and Meox2, which is contributes to tendon and soft connective tissue development (Acharya and Amit n.d.). Other regulators that have underexplored roles in IZ formation identified include Pitx1, Meox1, Deaf1, Tbx5, Jund, and Zeb1. Def1 has been reported to bind to Gdf5 and have a repressive effect on Gdf5 expression (Syddall et al. 2013). Tbx5 interacts with Fgf and Wnt in the limb bud to modulate limb and joint morphogenesis (Agarwal et al. 2003) (Rallis et al. 2003). Jund, along with Fos, forms a complex that directly regulates Wnt activity in the IZ (Kan and Tabin 2013). Zeb1 modulates TGFβ signaling, and when mutated leads to multiple joint fusions (Takagi et al. 1998). In summary, we have found that IZ formation is characterized by continuous chondrocyte-to-mesenchymal process that includes cells of the anlagen. Our analysis has revealed many previously implicated regulators of this process, as well as many novel candidate genes.

### Development of articular fibrous components

SC3, characterized by fibroblast-related genes and pathways, is distinct from the IZ-related SC2. As we did for SC1 and SC2, we performed a deeper analysis of SC3 by clustering it more finely into two sub-clusters. SC3_A was mainly comprised by E13.5 cells and had high levels of cell growth related genes whereas SC3_B was made up of E14.5 and E15.5 cells and had higher levels of fibroblast ECM related genes such as Postn and Col3a1 (**Fig 6A-B**). We confirmed the preferential expression of Col3a1 and Postn in the ligament, tendon and menisci by ISH (**Fig 5D** and **Fig 6C**). However, it was not clear whether intra-articular fibrous components (especially at E15.5) belonged to SC2 or SC3 based on the expression pattern of differential genes Postn, Col3a1, and Col2a1 (**Suppl Fig4A-E**). Nevertheless, the presence of Gdf5 at E15.5 articular surface and predominant expression in SC2 indicate intra-articular ligament cells were included in SC3 (**Suppl Fig4F**). Our data and analysis suggest that the Scx expressing cells of SC2 give rise to fibrochondrocytes, which contribute to the transitioning zone of articular cartilage, intra-articular ligament, and meniscus (**Suppl Fig4G**). In addition to Col3a1 and Col1a1, expression of Dcn and Tnmd were better able to differentiate SC3 from SC2 (**Suppl Fig4H,I**). Thus, Dcn^+^Tnmd^+^Scx^+^ SC3 refers to fibroblasts that contribute to fibrous tissue of joint.

**Figure 6:**
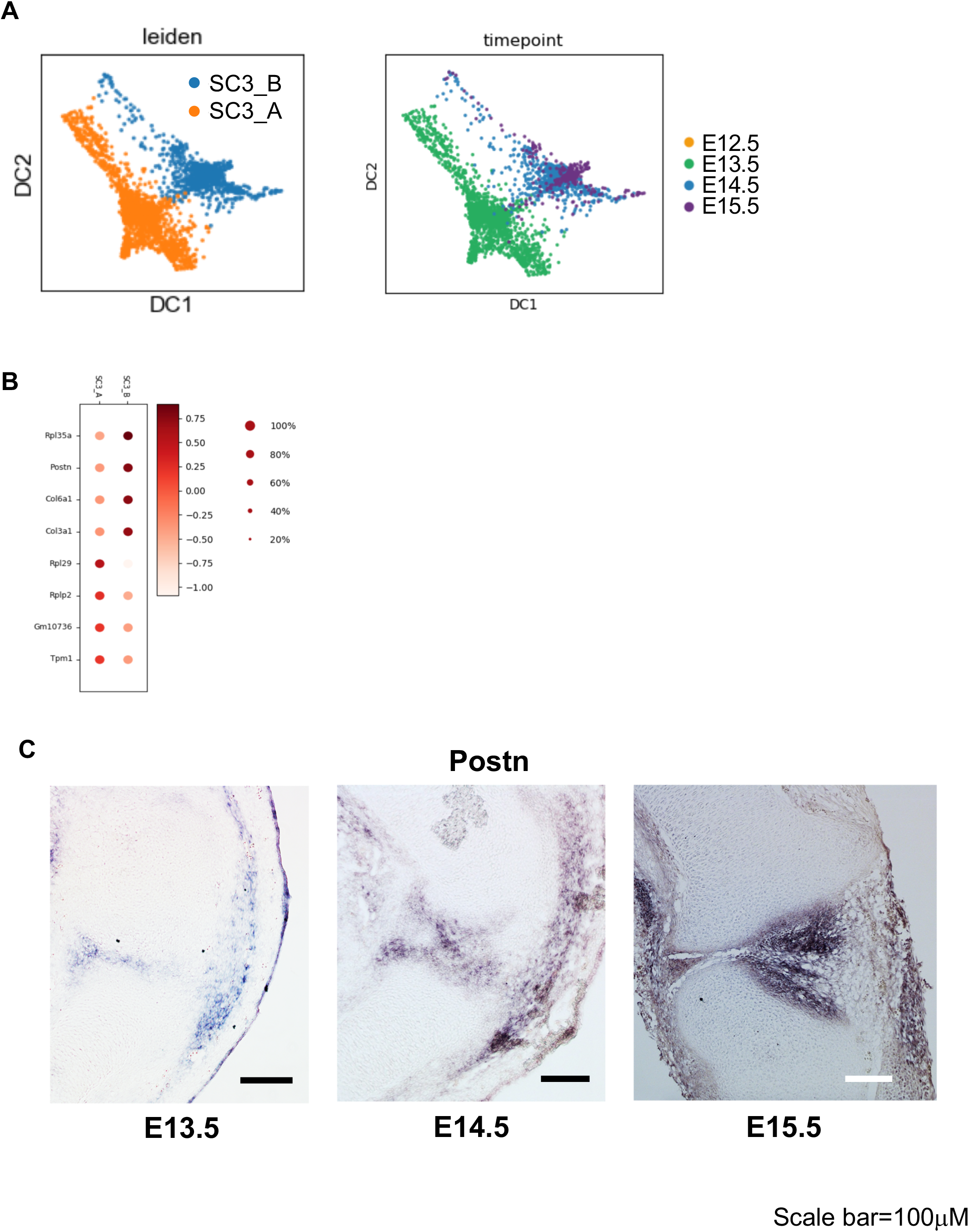
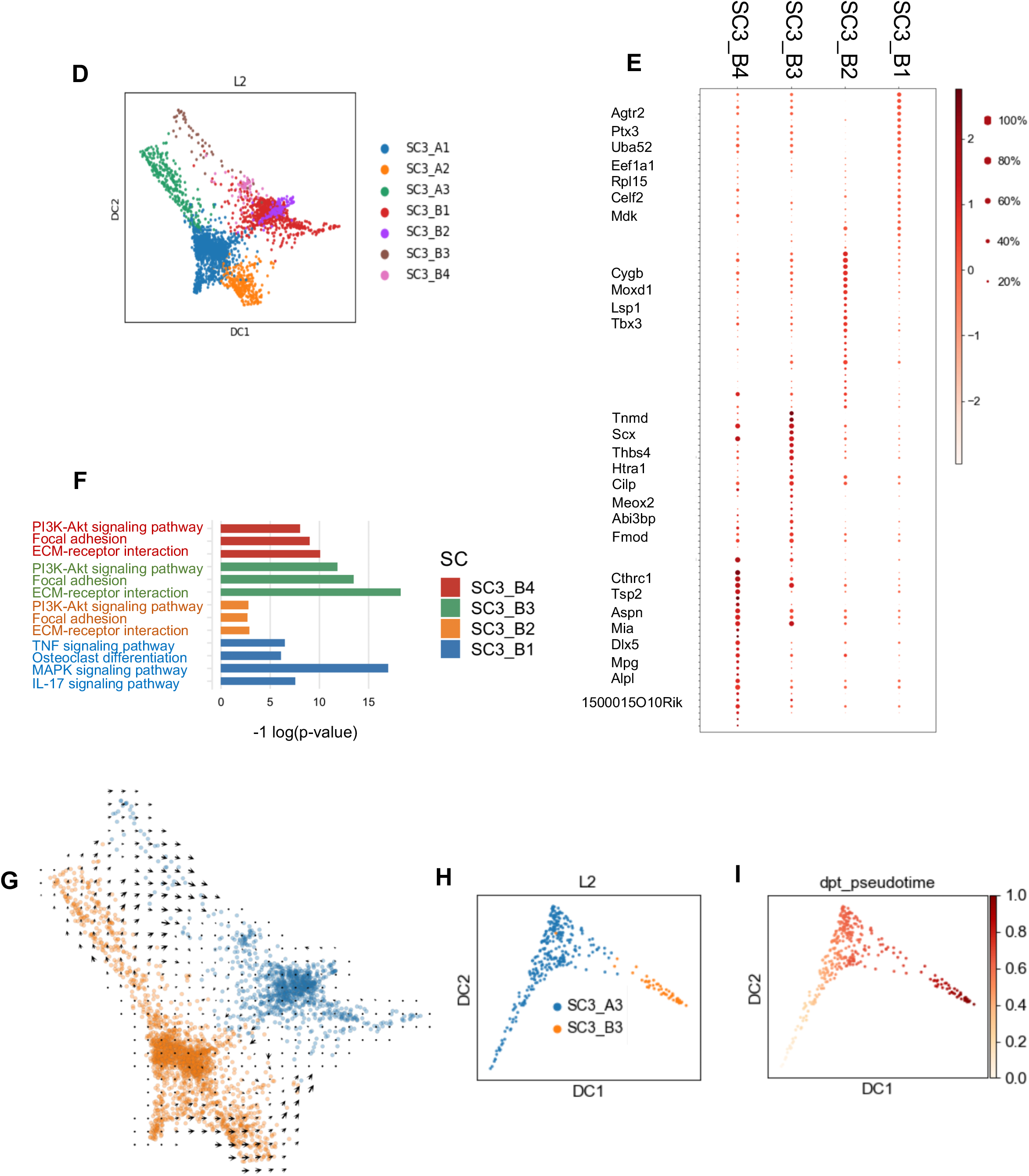
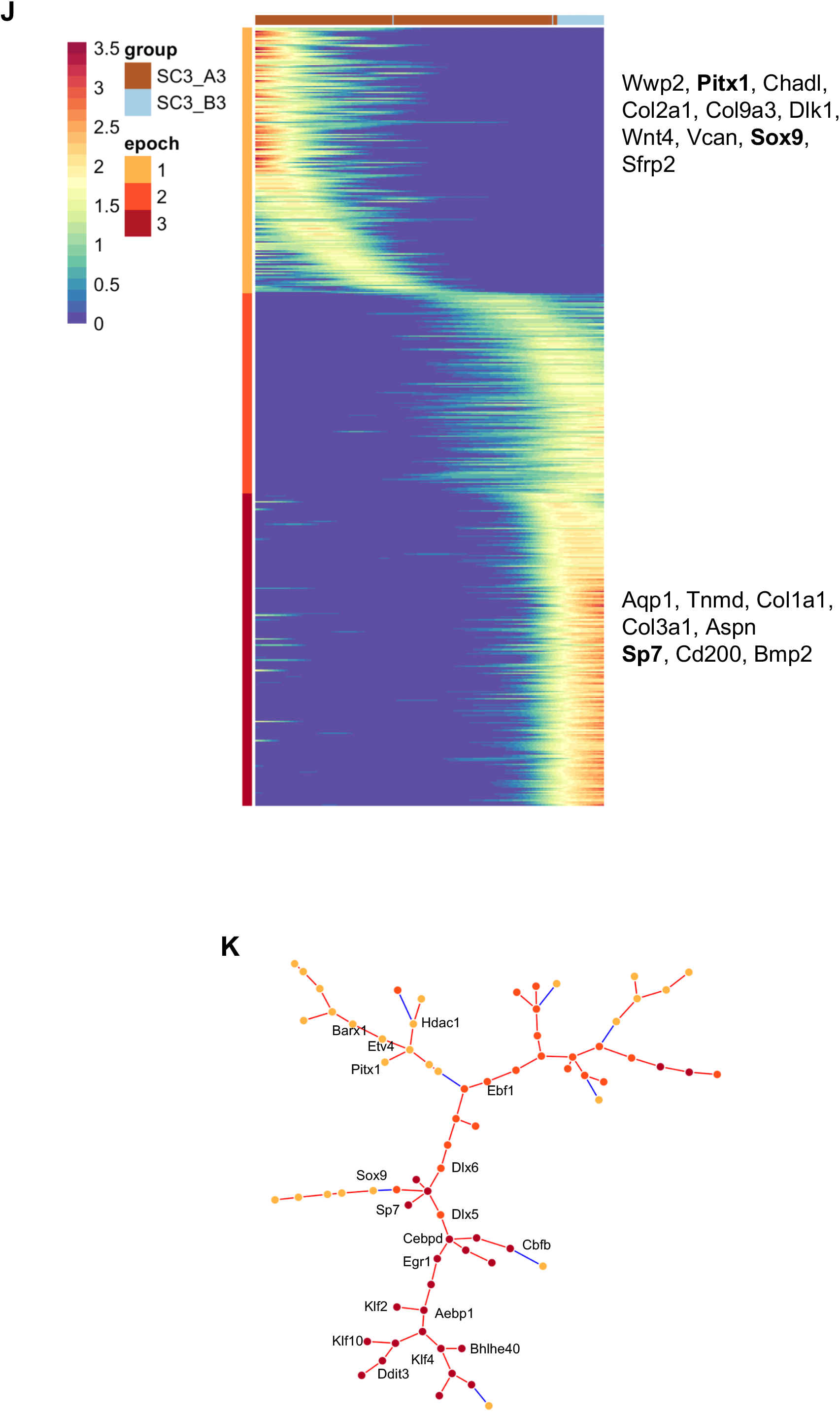
SC3 is composed of articular fibrous component cells. (A) Leiden clustering and diffusion map embedding SC3. (B) Dot plot expression of representative genes differentially expressed between SC3_A and SC3_B. (C) ISH detection for SC3_A representative genes. (D) Sub-clustering of SC3 by Leiden. (E) Dot plot expression of representative genes differentially expressed among 7 sub-clusters. (F) Enrichment analysis of SC3_B sub-clusters. (G) RNA Velocity analysis. Arrows indicate the predicted future state of SC3 cells, showing a minimal transition between SC3_A3 and SC3_B3. (H-I) Leiden clustering and diffusion map embedding SC3_A3 and SC3_B3, colored by groups (H) and pseudotime (I). (I) Epoch analysis identifies three Epochs of gene expression based on SC3_A3 and SC3_B3 populations. Selected genes from each Epoch are listed on right, and Epoch identified regulators are in bold. (J) Minimal spanning tree representation of Epoch-reconstructed gene regulatory network.

As SC3_A and SC3_B differed mainly by developmental stage, we sub-clustered each to search for more subtle differences in state or lineage, resulting in seven sub-groups (**Fig 6D**). We then annotated the likely cell type of each sub-cluster based on differential gene expression and enrichment analysis, as described below. SC3_B2 is likely to represent myotendinous junction site cells, as it has high levels of Tbx3, which is required for muscle attachment (Colasanto et al. 2016), in addition to other muscle-related genes including Lsp1, Cygb, and Moxd1 (Singh et al. 2014) (**Fig 6E**). Cells of SC3_B3 are likely to be tendon/ligament cells based on preferential expression of Tnmd and Scx (Sugimoto et al. 2013; Soeda et al. 2010) (Subramanian and Schilling 2015), as well as other tendon associated genes including Thbs4 (Havis et al. 2014) (Subramanian and Schilling 2014), Htra1 (Oka et al. 2004), Cilp (Caterson and Melrose 2018), Meox2 (Havis et al. 2014), Abi3bp (Zhang et al. 2014), and Fmod (Bi et al. 2007). SC3_B4 is likely to include synovial fibroblasts and fibrocartilage cells of the enthesis (Zelzer et al. 2014) based on the preferential expression of Cthrc1 and Tsp2, both of which are produced by synovial fibroblasts (Shekhani et al. 2016; Park et al. 2004) and chondrocyte-related genes Aspn, 1500015O10Rik, Mia, and Dlx5 (Ferrari and Kosher 2006). We hypothesize that SC3_B1 is comprised of synovial lining cells based on the enrichment of MAPK, IL-17, TNF signaling pathways, in contrast to the other SC3_B clusters, which were enriched in ECM-receptor interaction, Focal adhesion, and the PI3K-Akt signaling pathway (**Fig 6F**). Taken together, our data suggest that SC3_B subclusters represent cells of the myotendinous junction site (B2), tendon/ligament (B3), fibrocartilage cells of the synovium and enthesis (B4), and the synovial membrane (B1).

Next, we applied RNA velocity to infer lineage relationships among SC3 cells. This analysis detected velocity primarily between SC3_A3 and the tendon/ligament cluster SC3_B3 (**Fig 6G-I**). This result was consistent with the lineage annotation of SC_B subgroups we proposed above because there is little-to-no trajectory between the SC3_B sub-clusters. To perform Epoch analysis and reconstruct the GRN that contributes to this progenitor-to-tenocyte/ligamentocyte transition, we first performed diffusion based pseudotime analysis on SC3_A3 and SC3_B3 (**Fig 6H-J**). Our data and analysis are consistent with prior studies which reported that tendon/ligament progenitors lose Sox9 concomitant with Scx upregulation (Sugimoto et al. 2013; Soeda et al. 2010) and followed by Tnmd upregulation (Subramanian and Schilling 2015) (**Suppl Fig 4J**). Next, we used Epoch to identify genes temporally regulated along this pseudotime axis (**Fig 6I, Supp Table 4**). At the early stage, many chondrogenesis and IZ related genes were high including Wwp2, Pitx1 (Wang et al. 2018), Chadl, Col2a1, Col9a3, Dlk1(Chen et al. 2011), Wnt4, Vcan, Sox9, and Sfrp2. This suggested that the development of articular fibrous components, especially tendon/ligament starts from a progenitor population with some chondrocyte features. By the later stage, the mature tendon markers Aqp1, Tnmd, and the fibroblast ECM genes Col1a1, Col3a1, Aspn were upregulated (**Fig 6J**). Epoch analysis predicted that the early stage was regulated by Barx1, which has an inhibitory effect on chondrogenic initiation during joint development (Church et al. 2005). Other predicted regulators included Etv4, which is detected in muscle-tendon interface with high expression level and regulated by FGF signaling (Havis et al. 2016) and Hdac1, which was recently found to inhibit Scx expression in tendon progenitor cells (Zhang et al. 2018). Regulators of the middle stage included those previously associated with tendon development (e.g. Ebf1 --expressed in presumptive tendons surrounding chondrogenic condensation (Mella et al. 2004)) and other factors that have not previously been implicated in this process such as Dlx5, Dlx6, which are expressed in presumptive elbow joint and involved in osteogenesis (Ferrari and Kosher 2006; Lee et al. 2003). The predicted regulators of the final stage included TFs associated with inflammatory response: Cebpd, Egr1; osteogenesis: Sp7, Cbfb (Lien et al. 2007), and tendon development: Klf10 (McConnell and Yang 2010), Klf2, Klf4, Aebp1 (Blackburn et al. 2018), Ddit3 (Caterson and Melrose 2018), and Bhlhe40 (Peffers et al. 2015) (**Fig 6K**). In summary, the development of fibrous components of the synovial joint, particularly tendon/ligament, is characterized by the ordered loss of chondrogenic gene expression programs followed by the upregulation of tendon/ligament expression programs. Moreover, we found that the cells of different fibrous components can be distinguished by their transcriptional signatures.

### Nascent joint development

To better understand the potential lineage relationships of the superclusters, we applied RNA velocity to all GLE cells. We found that some chondroprogenitor SC1 cells were predicted to give rise to SC2_A1, and that the Osr1^+^Col3a1^+^ SC1_A cells were predicted to differentiate to SC3, consistent with their chondrogenic or mesenchymal features, respectively (**Fig 7A**). Synthesizing these results with prior analyses yielded the following summary of our data. Early GLE cells contained a CD9^+^ chondrogenic population and a PDGFRA^+^ mesenchymal population (**Fig 7B**). The chondrogenic progenitors gave rise to the IZ, which is comprised of Col2a1^+^Sox9^+^Col9a1^+^Gdf5^Low^ cells (SC2_A1, SC2_B2) and Sfrp2^+^Col3a1^+^Gdf5^high^ cells (SC2_A3, SC2_B1), which are likely to correspond to the outer and intermediate IZ, respectively. Our data supports the notion that some outer IZ serves as precursor for intermediate IZ. In addition, newly recruited Gdf5-expressing IZ cells with enrichment in Sfrp2, Htra1, Dkk3 (SC2_B3, B4) appear to develop to either more mature IZ cells or to fibrous cells of SC3. On the other hand, the mesenchymal progenitors of SC1_A differentiate to Col3a1^+^Postn^+^Dcn^+^Tnmd^+^ fibrous component cells, including ligament, tendon and synovium (SC3). Intriguingly, a group of Scx^+^Meox2^+^Meox1^+^Tbx5^-^ SC2 cells (SC2_A3, SC2_B1) was predicted to transit to SC3 (**Fig 7A**), suggesting that some of the fibrous components are specified from multiple origins, in this case from both the early SC1_A and the later, IZ SC2_B sub-cluster.

**Figure 7:**
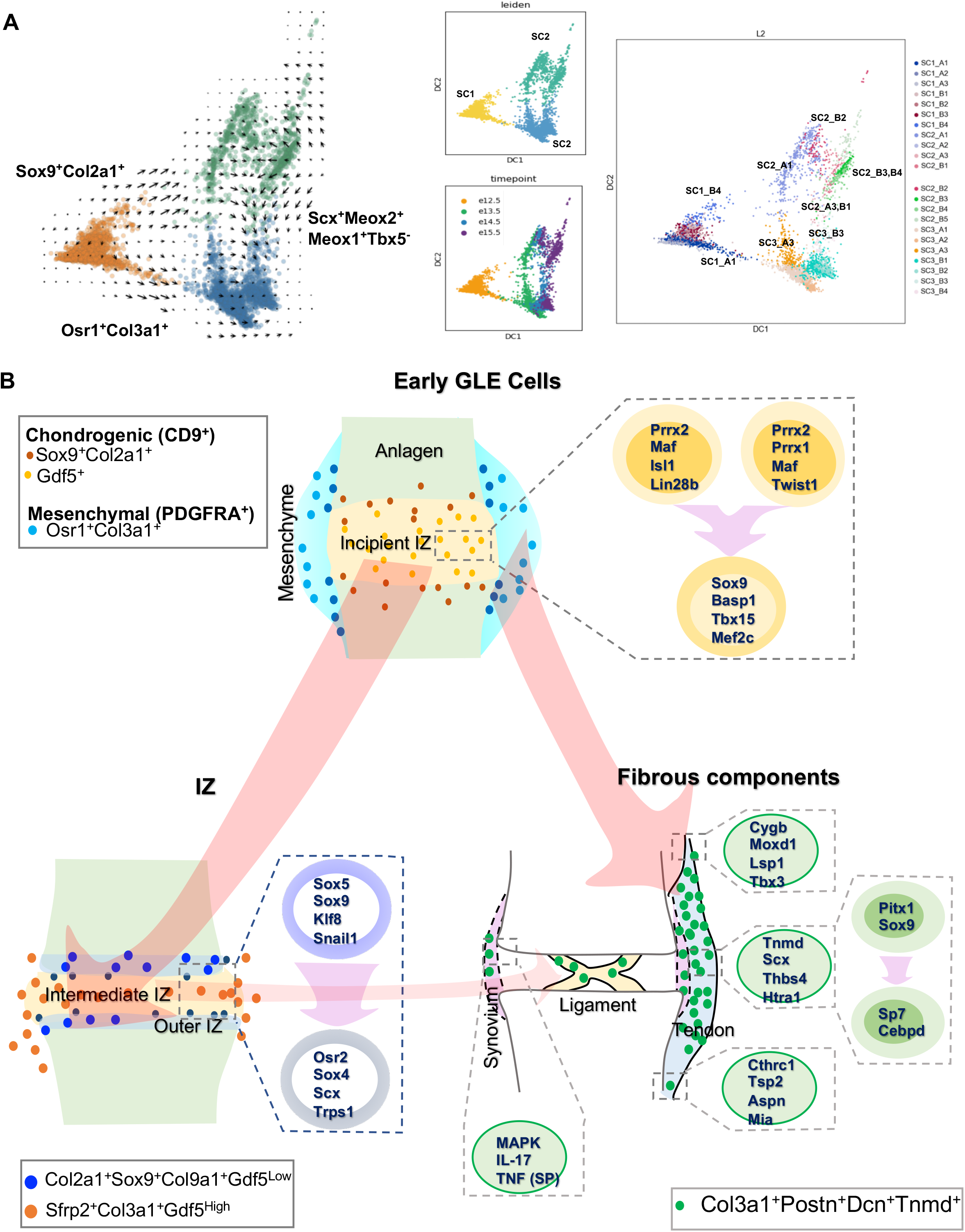
Nascent joint development. (A) RNA Velocity of 3 of SCs. (B) Cartoon of nascent joint development.

## Discussion

The synovial joint initiates from a thin layer of mesenchymal cells marked by Gdf5 expression. Through lineage tracing of Gdf5, it has become apparent that Gdf5-expressing IZ cells give rise to multiple joint lineages. However, the transcriptional programs that drive IZ formation and elaboration has remained underexplored. In this study, we applied scRNA-Seq to Gdf5 lineage cells during embryonic stages of synovial joint development to define the continuum of expression states that govern the process from interzone formation to joint cavitation. Here we have revealed the dynamic transcriptome changes and heterogeneity in GLE cells, we have inferred the lineage trajectories of subpopulations, and we have predicted the regulators of key developmental decisions. Several insights have emerged from our dataset and analyses that have implications for the field.

First, our results have revealed that GLE cells at E12.5 already exhibited a transcriptional heterogeneity, with one cluster tending towards a more mesenchymal state (SC1_A) and one cluster tending towards a more chondroprogenitor state (SC1_B). By prospectively isolating cells using markers that distinguished these clusters, we confirmed the *in vitro* lineage propensity of these cell populations. The degree of commitment of these cells *in vivo* remains to be determined. Second, we discovered further sub-structure within the SC1_B chondroprogenitors: one cluster (SC1_B1) preferentially expressed more proximal Meis2 and proximal/anterior Irx3, and one cluster (SC1_B2) preferentially expressed more distal (or zeugopod-associated) Hoxa11os. By RNA Velocity analysis, we predicted that both of these clusters were transitioning to a pre-IZ state marked by expression of Sfrp2, Vcan, Trps1, and Snail1, and our Epoch analysis revealed that the gene regulatory networks associated with these transitions were highly similar. This raises notion that the later complex architecture of the IZ and its derivatives are presaged by limb spatial patterning. Third, we found that between E13.5-E15.5 there is a continual transition of chondroprogenitors to an IZ state that is reminiscent of the pre-anlagen limb bud mesenchyme as exemplified by up-regulation of Prrx1. Fourth, our data support the idea that the fibrous joint components have dual origins. The Osr1^+^Col3a1^+^ mesenchymal progenitors detected in E12.5 (SC1_A) were predicted to transition to fibrous components of the tendon and fibrochondrocytes of the synovium, whereas cells of the putative intermediate IZ (SC2_ B3 and SC2_B4) were predicted to transition to Tnmd^+^Scx^+^ cells of the ligament/tendon cells (SC3_B3) (**Fig 7**).

We have made this data freely and easily accessible with a web application at http://www.cahanlab.org/resources/joint_ontogeny. We believe that this resource will aid the community in discovering additional transcriptional programs and in inferring cell interactions that underpin synovial joint development. Further, we anticipate that this data can be used to yield improved protocols for the derivation of synovial lineages from pluripotent stem cells(Oldershaw et al. 2010; Craft et al. 2015; Kawata et al. 2019; Yamashita et al. 2015) by, for example, using it to identify candidate signaling pathways or by using the expression data as a reference against which to compare engineered cells.

## STAR Methods

### Mice

*Gdf5-cre* (Sperm Cryorecovery via IVF, FVB/NJ background) mouse strain was obtained from the Jackson laboratory. B6.129X1-Gt(ROSA)26Sortm1(EYFP)Cos/J (RosaEYFP) was gifted by the lab of Prof. Xu Cao (Johns Hopkins University). Gdf5-cre::Rosa-EYFP mice were generated by crossing heterozygote Gdf5-cre strain with homozygote RosaEYFP strain. The genotype of the mice was determined by PCR analyses of genomic DNA isolated from mouse tails using the following primers: Gdf5-directed cre forward, 5’GCCTGCATTACCGGTCGATGCAACGA3’, and reverse, 5’GTGGCAGATGGCGCGGCAACACCATT3’ (protocol provided by Prof. David Kingsley, HHMI and Stanford University). Day 5 wild type refers to C57/BL10 mouse. All the protocols were approved by the institutional review board of Johns Hopkins University.

### Mice gender identification

We identified mouse gender by genotyping Sry Y gene using the primers: forward, 5’CTGGAAATCTACTGTGGTCTG3’, and reverse, 5’ACCAAGACCAGAGTTTCCAG3’.

### Cell isolation

Mice were kept in light-reversed room (light turns on at 10 pm and turns off at 10 am). Timing was determined by putting one male mouse and two female mice in the same cage at 9 am and separating them at 4 pm on the same day. We count that midnight as E0.5 stage. On E12.5, E13.5, E14.5 and E15.5, the pregnant mice were sacrificed by CO_2_ at 3 pm. The cells were isolated using the protocol (Primary culture and phenotyping of murine Chondrocytes) with modification: The embryos (usually n=6-8) were rinsed three times in PBS on ice. Two presumptive joint part from hind limb between presumptive ankle and hip of each individual embryo were disassociated in a single 3 cm dish (Figure 1A) and incubated in digestion solution I (3 mg / mL collagenase D, DMEM high glucose culture medium, serum free) for 45 min at 37 °C, and then in digestion solution II (1 mg / mL collagenase D, DMEM high glucose culture medium, serum free) for 3 hrs (one embryo per dish) at 37 °C. During the period of incubation, the mice gender was identified by genotyping and only male samples were chosen for further processing. The tissues with medium were gently pipetted to disperse cell aggregates and filtered through 40 μm cell strainer, then centrifuged for 10 min at 400 g. The pellet was suspended with 0.4% BSA in PBS.

### Cell fractionation

All cells were fractionated by fluorescence-activated cell sorting (FACS). A MoFlo XDP sorter (Beckman Coulter, Miami, FL. USA) was used to collect YFP^+^ cells, and Propidium Iodide was used to exclude dead cells.

### Single cell RNA sequencing

GemCode™ Single Cell platform (10X Genomics) was used to determine the transcriptomes of single cells (Zheng et al. 2017). Cells at 1000 / μl were obtained after sorting and placed on ice. Each time point, one sample was selected and profiled based on the viability and amount. A total of 6000 cells were loaded each time, followed by GEM-RT reaction, and cDNA amplification. Single cell libraries were constructed by attaching P7 and P5 primer sites and sample index to the cDNA. Single cell RNA sequencing was performed on Illumina NextSeq 500 and HiSeq 2500 to a depth ranging from 347 to 489 million reads per sample.

### Analysis and visualization of scRNA seq data

CellRanger (version 2.0.0) was used to perform the original processing of single cell sequencing reads, aligning them to the mm10 reference genome. We used the command line interface of Velocyto, version 1.7.3, to count reads and attribute them as spliced, un-spliced, or ambiguous (La Manno et al. 2018). The resulting loom files for each sample were then concatenated and subjected to quality control processing, normalization, estimation of cell cycle phase, clustering, and differential gene expression analysis using Scanpy 1.4.3 (Wolf et al. 2018). Specifically, we excluded cells in which mitochondrial gene content exceeded 5% of the total reads or cells in with fewer than 501 unique genes detected. Then, we excluded genes that were detected in fewer than 10 cells, resulting in a data set of 10,124 cells and 16,352 genes. Then, we performed an initial normalization on a per cell basis followed by log transformation, and scaling. We scored the phases of cell cycle using cell cycle-associated genes as previously described (Satija et al. 2015). Then we identified the genes that were most variably expressed across the whole data set, and within each timepoint, resulting in 3,593 genes. We performed PCA and inspected the variance ratio plots to determine the ‘elbow’, or number of PCs that account for most of the total variation in the data. We generated a graph of cell neighbors using diffusion maps (Coifman et al. 2005), and then we performed Leiden clustering (Traag et al. 2019), which we visualized with a UMAP embedding (McInnes and Healy 2018). We also analyzed this with SingleCellNet (Tan and Cahan 2019), which had been trained using the Tabula Muris data set (Tabula Muris Consortium et al. 2018). We removed cells in clusters that were classified by SingleCellNet as ‘blood’, ‘erythroblast’, ‘endothelial’. We also removed cells in clusters that we identified as likely to be myoblast based on high levels of Myod1 and other muscle-specific genes, melanocyte (based on Pmel expression), and neural crest (based on Sox10 expression). Then, we repeated the pre-processing and analysis pipeline on the remaining 8,378 genes. We noted that two clusters, primarily from E12.5 and E13.5, were predicted to be in G2M phase; we removed these cells from further analysis. Finally, we removed cells in a cluster that we determined by ISH to consist mainly of dermis cells, resulting in final data set of 6,202 cells and 16,352 genes. Super-clusters and all sub-clusters were identified by following the same pipeline as described above, except that the analysis was limited to the corresponding set of cells. For example, the superclusters were identified by first finding the genes that vary across both all cells, and within each time point. Then, a neighborhood graph was determined using the principal components (the number of which was decided by examining the variation ratio plot), followed by Leiden clustering, and visualized by UMAP embedding, and, for some subsets of data, diffusion map embedding. Differentially expressed genes were identified using the Scanpy rank_genes_groups function. Gene set enrichment analysis was performed using GSEAPY (https://github.com/zqfang/GSEApy), a Python interface to enrichR (Chen et al. 2013; Kuleshov et al. 2016). The analysis pipeline of Velocyto was applied to data subsets as mentioned in the main text. We used the Velcoyto results to manually assign roots for diffusion map pseudotime analysis. The results of pseudotime were imported into Epoch for gene regulatory network reconstruction (manuscript in preparation).

### Histochemistry, immunohistochemistry, and histomorphometry

The specimens were fixed in 10% buffered formalin for 6-24 hrs at RT. D5 joints were decalcified in 10% ethylenediaminetetraacetic acid (EDTA) in PBS (pH 7.4) for 3 days at 4°C, washed with distilled water and equilibrated in 30% sucrose in PBS at 4°C overnight, then mounted in O.C.T and frozen at −80°C. Ten-micrometer-thick coronal-oriented or sagittal-oriented sections were performed by cryostat.

We performed Trichrome staining according toTrichrome Stain (Connective Tissue Stain) Kit protocol.

Immunostaining was performed using a standard protocol. Sections were incubated with primary antibodies to mouse GFP (1:200), TNMD (1:100), SOX9 (1:500), THY1 (1:100) in Antibody Diluent, at 4ū°C overnight followed with three 5 min washes in TBST. The slides were then incubated with secondary antibodies conjugated with fluorescence at room temperature for 1 ūh while avoiding light followed with three 5 min washes in TBST and nuclear stained with mounting medium containing DAPI. Images were captured by Nikon EcLipse Ti-S, DS-U3 and DS-Qi2. See **Suppl Table 5**.

### *In situ* hybridization

See **Suppl Table 6** for the information of oligonucleotides used for templates for antisense RNA probes. The T7 and SP6 primer sequence were added to 5- and 3-prime end, respectively. SP6 RNA polymerase was used for probe transcription. Probes were synthesized with digoxygenin-labeled UTP and hybridized at 68 °C overnight. Results were visualized by Alkaline phosphatase-conjugated anti-digoxygenin antibody and BCIP/NBT substrates.

### FACS for prospective isolation

E12.5 embryonic hind limb cells or Day 5 knee joint cells were isolated as described in Cell isolation. After filtered through 40 μm cell strainer, cells were suspended in autoMACS rinsing solution at 1 million per mL. After spin down, E12.5 cells were then stained with PDGFRA (1 μg per 10 million cells) and CD9 (1 μg per 5 million cells) in 100 uL autoMACS rinsing solution in dark for 30 min followed by two times washes with autoMACS rinsing solution. Cells were resuspended in autoMACS rinsing solution. A negative control without staining was used to setup gate. The following two E12.5 populations were collected at the same time: YFP^+^PDGFRA^+^ population, YFP^+^PDGFRA^-^CD9^+^ population. Day 5 four populations were collected based on four evenly distributed cell samples.

---

**Supplemental Table 1**

TF score for SC1_B Path 1. TF: Transcription factor; epoch: 1=Early stage; 2=Middle stage; 3=Later stage; weightMean: mean association strength of TF and targets genes; ntargets: Predicted number of target genes; peakTime: the pseudotime at which gene expression is highest.

**Supplemental Table 2**

TF score for SC1_B Path 2.

**Supplemental Table 3**

TF score for SC2 Path.

**Supplemental Table 4**

TF score for SC3_A3 to B3 Path.

## Acknowledgements

This work was supported by the National Institutes of Health under grant R35GM124725 to PC and by the Maryland Stem Cell Research Fund 2017-MSCRFF-3910 (Award ID: 90074850) to Qin Bian. Jordan Wilson was supported by NIH R25GM109441. This work was made possible by support from the Johns Hopkins Medicine Discovery Fund.

