## Supplemental figures and tables for "A single cell transcriptional atlas of early synovial joint development"

**Supplemental Information**

### Supplemental Figure 1

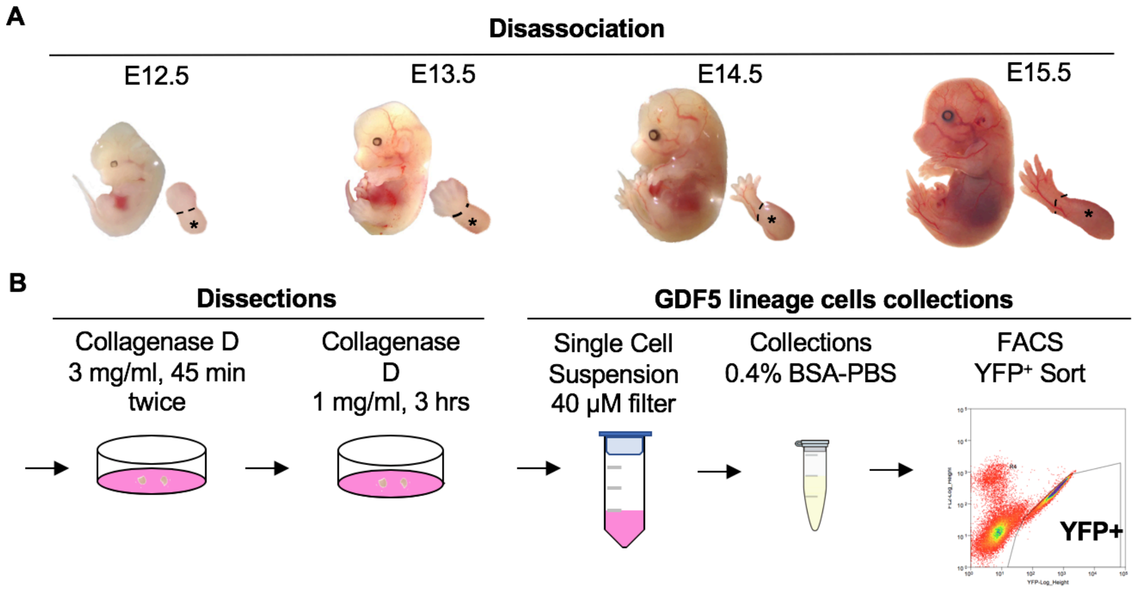


**Supplemental Figure 1**: (A) Developmental stage and region of hind limb dissected for cell isolation. (B) Schematic of single cell dissociation and YFP$^{+}$cell isolation.

**Supplemental Figure 2**

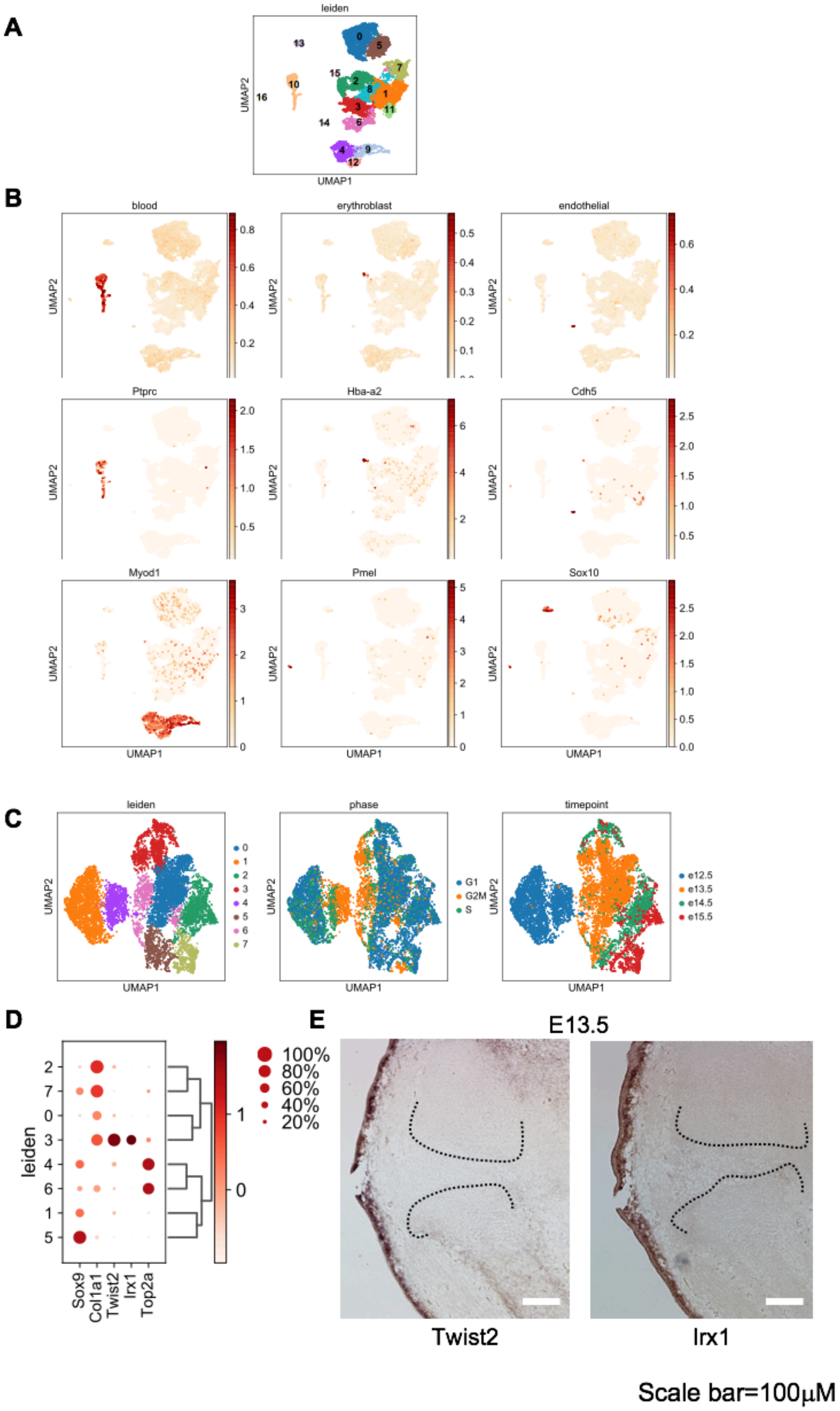


**Supplemental Figure 2**: Initial clustering and identification of non-joint cells and clusters of cells defined by stage of cell cycle.

### Supplementary Figure 3

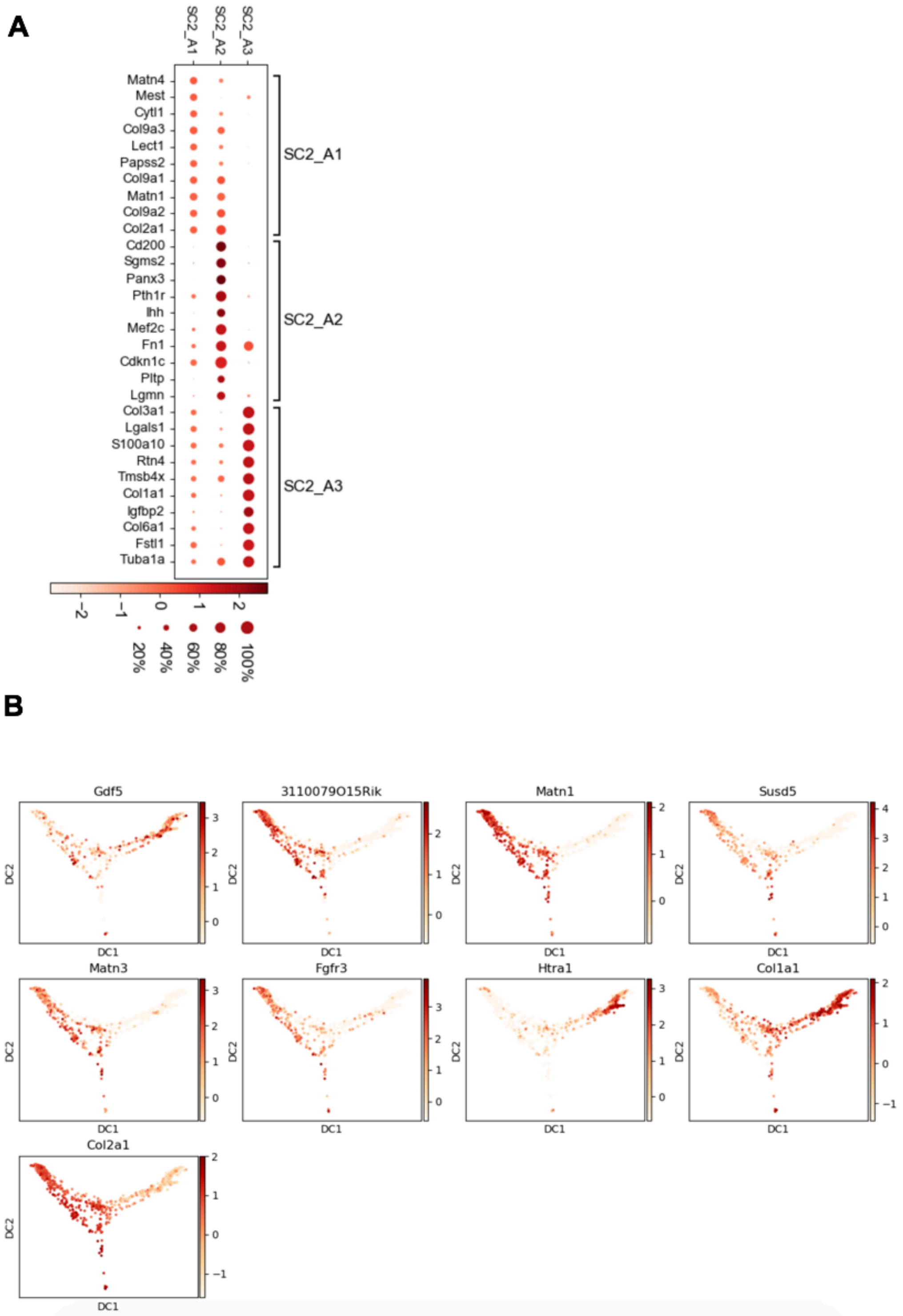


**Supplemental Figure 3**: Expression distribution of outer IZ and intermediate IZ representative genes of SC2.

### Supplementary Figure 4

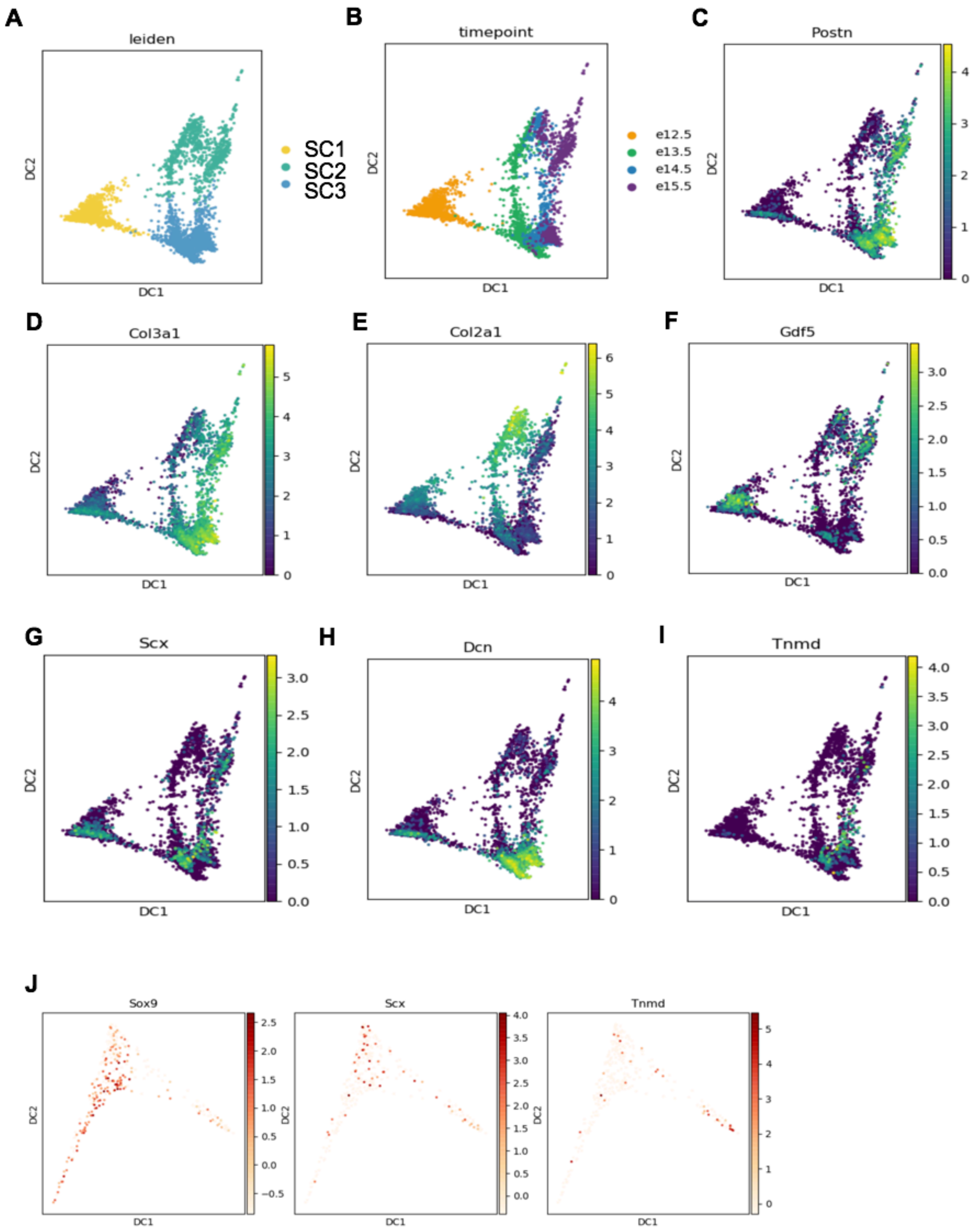


**Supplemental Figure 4**: (A, B) Leiden clustering of SCs, colored by groups (A) and timepoints (B). (C-I) Expression distribution of representative genes on 3 of SCs. (J) Typical tendon/ligament developmental marker genes expression on SC3_A3 and SC3_B3 arranged by pseudotime.

### Supplemental Table 5

#### Reagent Source

| REAGENT | SOURCE | IDENTIFIER |
|---|---|---|
| **Antibodies** | | |
| **Anit-GFP antibody (Rabbit polyclonal to GFP)** | Abcam | Ab6556 |
| **Anti-TNMD antibody (Rabbit polyclonal to TNMD)** | Abcam | Ab203676 |
| **Anti-SOX9 antibody (Rabbit monclonal to SOX9)** | Abcam | Ab185966 |
| **Anti-THY1 antibody (Rabbit monclonal to THY1)** | Abcam | Ab3105 |
| **Goat anti-rabbit IgG H&L(Alexa Fluor 488)** | Abcam | Ab150077 |
| **Goat anti-rabbit IgG secondary antibody, Cy3 conjugate** | Boster | BA1032 |
| **CD140a (PDGFRA) monoclonal antibody, APC** | eBioscience | 17-1401-81 |
| **CD9 monoclonal antibody, eFluor 450** | eBioscience | 48-0091-82 |
| **Chemicals, enzymes, recombinant proteins** | | |
| **Antibody Diluent** | Agilent Dako | S080981-2 |
| **Vectashield Antifade Mounting Medium with DAPI** | Vector Laboratories | H-1200 |
| **Trichrome Stain (Connective Tissue Stain) kit** | Abcam | ab150686 |
| **Propidium Iodide** | Sigma | P4864 |
| **GenElute™ Mammalian Total RNA Miniprep Kit** | Sigma Aldrich | RTN70 |
| **autoMACS ™ Rinsing Solution** | Miltenyi Biotec | 130-091-222 |
| **SuperScript™ III Reverse Transcriptase** | Invitrogen | 18080044 |
| **Collagenase D** | Roche | 11088858001 |
| **Extracta™DNA Prep for PCR** | QuantaBio | 95091-025 |
| **DreamTaq Green PCR Master Mix (2X)** | Thermo Scientific™ | K1081 |
| **iTaq Universal SYBR Green Supermix** | BIO-RAD Laboratories | 1725121 |
| **RNase A** | Roche | 10109142001 |

| | | |
|---|---|---|
| **Proteinase K** | Sigma-Aldrich | P2308-25MG |
| **DIG RNA Labeling Mix** | Roche | 11277073910 |
| **SP6 RNA Polymerase** | Millipore Sigma | 10810274001 |
| **BCIP 5-bromo-4-chloro-3-indolyl-phosphate, 4-toluidine salt** | Millipore Sigma | 11585002001 |
| **NBT Substrate powder(nitro-blue tetrazolium chloride)** | Thermo ScientificTM | 34035 |
| **DNA, MB-grade from fish sperm** | Millipore Sigma | 11467140001 |
| **tRNA from brewer's yeast** | Millipore Sigma | 10109525001 |
| **MEM α, no nucleosides** | Gibco™ | 12561056 |
| **Smooth Muscle Cell Differentiation Medium** | Sigma-Aldrich | 311D-250 |
| **Synoviocyte Growth Medium** | Sigma-Aldrich | 415F-500 |
| **Horse Serum** | Gibco™ | 16050114 |
| **Recombinant Mouse BMP-6 Protein** | R&D | 6325-BM-020 |
| **Chicken Embryo Extract Powder** | Gemini | 100-163P |
| **TGF Beta 3** | Lonza | PT-4124 |

### Supplemental Table 6

Oligonucleotides for PCR amplification of templates for antisense RNA probes

| Gene Symbol | Forward Primer | Reverse Primer | Probe size (bp) |
|---|---|---|---|
| ***Osr1*** | AAGCGTCAGAAGTCTAGTTCG | GCTTCTTTTCTGGGGATAGCTT | 634 |
| ***Col2a1*** | GTCCTACTGGAGTGACTGGTCC | CCAGATTCTCCTTTGTCACCTC | 738 |
| ***Irx1*** | ATGTCCTTCCCGCAG | TCAGGCAGACGGGAG | 1444 |
| ***Sfrp2*** | AGCAACTGCAAGCCCATC | ATGGAGAGAAGCCACCCC | 803 |
| ***Twist2*** | CGCCAGGTACATAGACTTCCTC | GTAAAGAACAGGAGTATGCGGG | 675 |
| ***Col9a1*** | AGAGGCCAGATTGATGCG | CATCAAATCCCCGAGCAC | 843 |
| ***Postn*** | TTTAGAGCAGCCGCCATC | CTGCAGCTTCAAGGAGGC | 811 |
| ***Col3a1*** | CTCAGGGTATCAAGGGTGAAAG | AGACTTTTCACCTCCAACTCCA | 739 |
| ***Sox9*** | ATGAATCTCCTGGACCCC | TCAGGGTCTGGTGAGCTGTG | 1532 |
| ***Gdf5*** | GCCTTGTTCCTAGTGTTTGGTC | CAGCCCCTGTAATGAACATCTC | 899 |
